# Temporal resource continuity increases predator abundance in a metapopulation model: insights for conservation and biocontrol

**DOI:** 10.1101/2020.03.25.006882

**Authors:** Brian J. Spiesman, Benjamin Iuliano, Claudio Gratton

## Abstract

The amount of habitat in a landscape is an important metric for evaluating the effects of land cover and land use on biodiversity and ecosystem services, yet it fails to capture complex temporal dimensions of resource availability that could be consequential for species population dynamics. If ephemeral resources across multiple habitat patches are synchronously available, resource gaps could be detrimental to population growth. In contrast, asynchronously available resources create a mosaic of temporally complementary resources that mobile organisms can track across the landscape. Knowledge is especially lacking on the relevance of temporal complementation for tri-trophic interactions and biological pest control. Here we use a spatially-explicit predator-prey metapopulation model to test the effect of different spatiotemporal resource patterns on insect predators and their prey. We examined prey and predator responses in model landscapes that varied in both the amount and temporal variability of basal vegetation resources. Further, we examined cases where prey comprised either a single generalist species or two specialist species that use different resources available either early or late in the growing season. We found that predators and generalist prey benefitted from lower temporal variance of basal resources, which increased both of their landscape-scale abundances. However, increasing the amount of basal resources also increased the variability of generalist prey populations, resulting in a negative correlation between basal resource amount and predator abundance. Specialist prey, on the other hand, did not benefit from less temporally variable in basal resources, since they were restricted by habitat type while also suffering greater predation. Predators feeding on specialists achieved greater prey suppression in landscapes with less temporally variable resources. Our simulations demonstrate the joint importance of landscape-scale temporal dynamics of resources and resource amount in understanding how landscape heterogeneity influences biodiversity and ecosystem services such as the biological control of agricultural pests.

## Introduction

The abundance and diversity of species in a landscape generally increases with greater habitat area (Fahrig 2003), presumably because larger areas can provide more limiting resources to consumers. This is a central principle of biodiversity conservation and the conservation of beneficial species that provide ecosystem services in working landscapes. For example, increasing non-crop habitat surrounding agricultural fields can promote insect abundance, diversity, pollination, and natural pest control (Dainese et al. 2019, Kennedy et al. 2013, Chaplin-Kramer et al. 2011, Ricketts et al. 2009, but see Karp et al. 2018). Yet a facile understanding of habitat vs. non-habitat or cropland vs. natural area ignores the substantial heterogeneity that exists within land use categories (Vasseur et al. 2013) as well as the complex temporal dynamics of mobile consumers and their resources across the landscape (Cohen & Crowder 2017, Schellhorn et al. 2015).

“Landscape complementation” describes the processes by which organisms acquire key resources by traversing multiple patch types in a landscape (Dunning et al. 1992). This process may be especially important in agriculturally-dominated landscapes, where temporal gaps or bottlenecks in food availability brought about by landscape simplification—e.g. increasing annual cropland area or decreasing crop diversity—may negatively affect beneficial species that require nourishment continuously throughout the growing season (Schellhorn et al. 2015). Differences between landscapes in the composition of habitats can give rise to differential temporal patterns of resource availability for mobile consumers. In homogenous landscapes, temporal gaps could result if ephemeral resources emerge synchronously across different patches (Fig. 1A and C) and then disappear for the rest of the growing season (e.g., before planting or after harvesting in a monocropped landscape). Although mobile organisms could travel within a landscape to access potential food resources in different locations, there may be temporal gaps at critical points in the organism’s life cycle which could affect growth, survival, and reproduction. On the other hand, food resources in different habitat patches could emerge asynchronously (Fig. 1B and D). Such asynchrony creates an ensemble of temporally complementary patches that minimizes resource gaps and, for mobile species, results in continuous resource provisioning at a landscape scale over the course of a season (e.g., in a landscape of phenologically divergent annual and perennial crops). We refer to this process “temporal resource complementation.”

**Figure 1.**
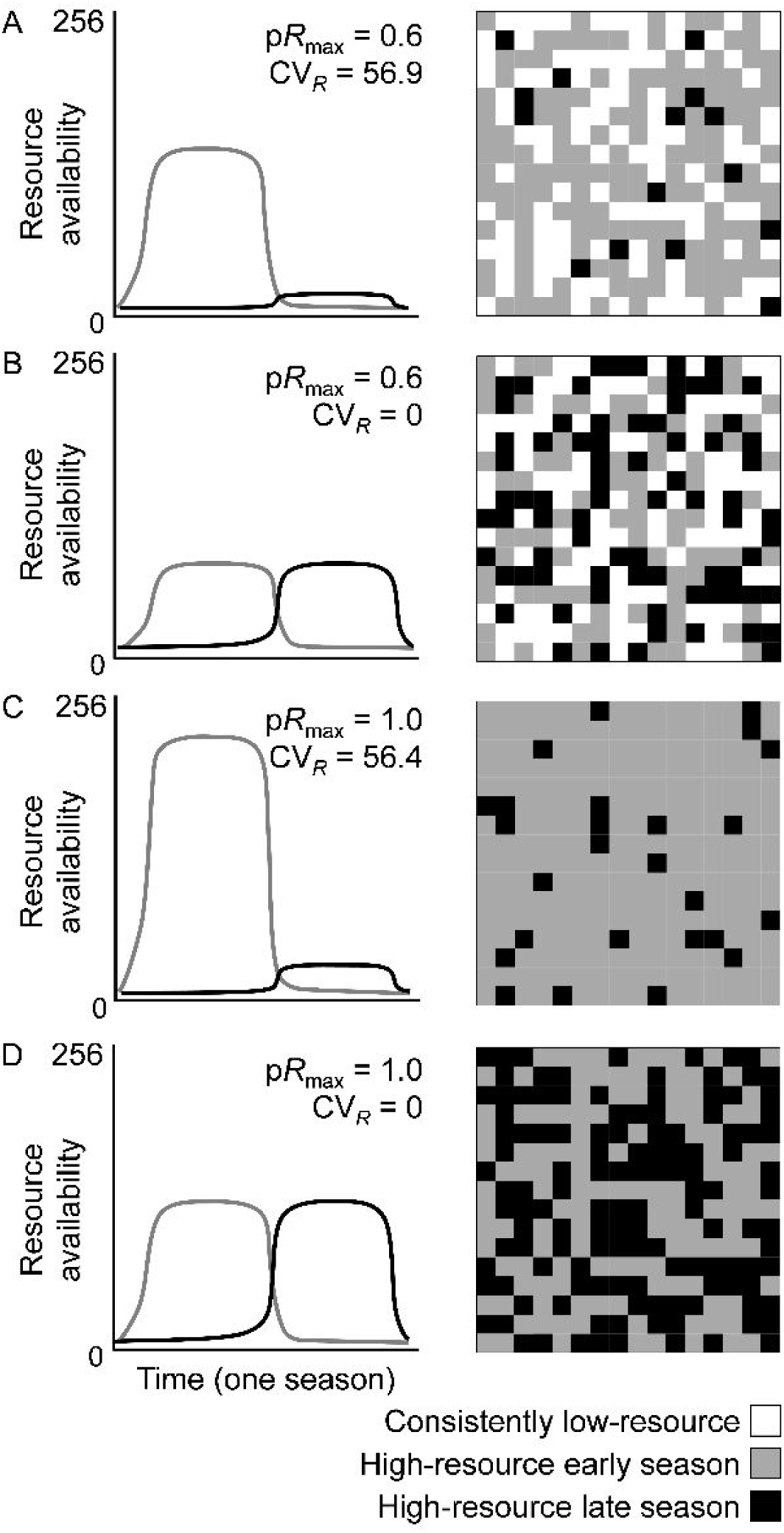
Conceptual diagrams (left) and corresponding model landscapes (right) for landscapes with different proportions of high-resource habitat (p*R*_max_) and temporal variability (*CV*_*R*_). Gray lines represent the amount of resources provided by habitat A (gray patches). Habitat A provides abundant basal resources for the first half of the season (i.e., *K* = 100 for prey *G* [generalist] and *S1* [specialist]), but few basal resources during the second half of the season (i.e., *K* = 10 for prey *G* and *S1*). Similarly, black lines represent the amount of resources provided by habitat B (black patches). Habitat B provides abundant resources (i.e., *K* = 100 for prey *G* and *S2*) during the second half of the season and few resources (i.e., *K* = 10 for prey *G* and *S2*) during the first half. White patches provide few basal resources year-round (i.e, *K* = 10 for all prey). In these model landscapes, there is no patch type providing abundant resources over the entire growing season. In panels A and B, both landscapes provide abundant basal resources in 60% of the patches over the full course of the season (p*R*_max_ = 0.6) but the temporal variance in resource availability is relatively high in panel A (*CV*_*R*_ = 56.9, i.e., basal resources are abundant early [90% of available resources] but rare late in the season [10% of available resources]) and low in panel B (*CV*_*R*_ = 0, 50% of available resources available in each half of the season). In panels C and D, both landscapes provide high-resource resources in 100% of the patches (p*R*_max_ = 1.0), but only for a portion of the growing season. The temporal variance in resource availability is relatively high in panel C (*CV*_*R*_ = 56.4 [90% of available resources present early, 10% present late]) and low in panel D (*CV*_*R*_ = 0 [50% of available resources present in each half of the season]).

The value of temporal resource complementation has received empirical support in beneficial insects such as wild bees, which can benefit from the existence of temporally asynchronous floral resources in heterogeneous landscapes (Mallinger et al. 2016, Rundlöf et al. 2014, Mandelik 2012). However, much less is known about how temporal resource complementation affects predatory insects, the dynamics of tritrophic interactions, and the consequences for conservation biological control (*sensu* Begg et al. 2017). A population of mobile predators may track ephemeral prey resources across a landscape as they emerge in one location, decline, and then emerge in another. Yet a predator population’s ability to track herbivorous prey depends on the prey’s ability to track their own resources, constituted by the vegetation present in the landscape (hereafter “basal resources”). Habitat-generalist prey may benefit more from temporal resource complementation than habitat-specialists if mobile generalists can utilize multiple basal resource types that emerge in different locations throughout the season. Generalist prey may thus support larger predator populations than would specialist prey. However, it is not clear whether (1) an increase in predator population size would enhance generalist prey suppression or (2) lower predator density could maintain suppression of specialist prey. This latter scenario might be especially relevant to biological control in agroecosystems, where many pests, such as aphids, specialize on particular crop types (e.g., grains or legumes) but are frequently consumed by generalist predators such as lady beetles (Obrycki & Kring 1998).

Due to the difficulty of conducting empirical studies rigorously investigating the spatio-temporal dynamics of insect populations across multiple generations, modeling approaches are frequently employed (e.g. Levins 1969, Ives & Settle 1997, Bianchi & Van der Werf 2003). Here we use a spatially explicit predator-prey metapopulation model (Taylor 1990) to examine how patchy landscapes differing in the amount and continuity of basal resources affect the population dynamics of mobile arthropod herbivores (specialist or generalist prey) and, in turn, their arthropod natural enemies (generalist predators). Model landscapes are comprised of patches where ephemeral resources emerge either early or late in a growing season. This allows us to simulate different amounts of basal resources within and across a growing season as well as different levels of resource continuity, which we measure as the landscape-scale temporal variance of basal resources across time (hereafter “temporal variance”).

We predicted that landscapes with a greater total amount of basal resources would support larger prey and predator populations, and that greater temporal continuity (i.e., lower temporal variance and greater temporal complementation) would increase population sizes due to less frequent landscape-scale temporal resource gaps (Schellhorn 2015). We compare the response of a predator-prey system with a single generalist prey species that can use both early and late-season resource patches to that of a system with two habitat specialists, each of which specializes on resources in different habitats that emerge either early or late in the growing season. We predicted that a system with one generalist prey species would support larger predator populations than a system with two specialist prey species.

## Materials & Methods

### Model landscapes

Predator and prey population dynamics were modeled in each cell of a 16 × 16-cell lattice, representing a square landscape with 256 patches (e.g., Fig. 1). Patches vary in their capacity to provision basal resources, which determines the carrying capacity (*K*) of prey populations. The habitat in each patch *x* at location *i* can switch mid-season between a high-resource state (*x*_*i*_ = 1) that offers relatively abundant basal food resources for prey populations, or a low-resource state (*x*_*i*_ = 0) that offers relatively few resources for prey. We thus defined three states that a patch could assume throughout a growing season: 1) early-season high-resource (e.g. early-season crop), 2) late-season high-resource (e.g. late-season crop), or 3) continuously low-resource (e.g. highly disturbed or developed area). Because our focus in this study is investigating the potential role of cropland as habitat and simulating dynamics of predators and prey in managed agricultural landscapes, we intentionally exclude a fourth potential resource state, that is one of continuously high-resources (e.g., diverse natural habitats that have resources early and late in the season). By varying the number and location of high-resource patches in each half of the growing season, we can generate model landscapes with different combinations of basal resource amount and temporal variance (Fig. 1). We quantify the total, or season-long, amount of resources *R* in a landscape as the sum of resource state values in each half (*A* and *B*) of the growing season:

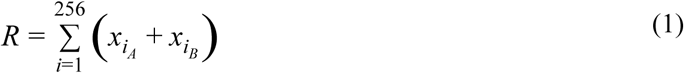

Thus, *R* can vary between 0 and 256 in our model landscapes. Landscape-scale temporal variance in basal resource amount is the variance in sum of patch values between the first and second half of the season:

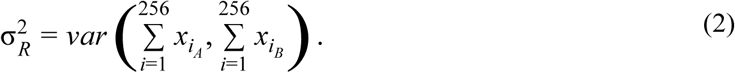

Because *R* varies among model landscapes, we use the coefficient of variation of (*CV*_*R*_) to standardize for the level of *R*, where

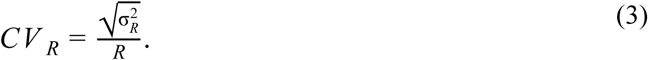

Thus, for a given level of *R*, high *CV*_*R*_ values indicate landscapes where resources are synchronously available and concentrated in only half of the growing season, which creates a “gap,” or period of low resource availability, during the other half of the season (Fig. 1A and 1C). In contrast, low *CV*_*R*_ values indicate landscapes where resources are asynchronously available and more evenly distributed throughout the growing season (Fig. 1B and 1D).

### Predator-prey metapopulation model

We modeled the spatially-explicit metapopulation dynamics of a generalist predator (e.g., a predatory arthropod) and its generalist or specialist prey (e.g., herbivorous insect), which reproduce and disperse on the 256-patch model landscapes. In short, subpopulations of a predator species, *P*, within each patch consume prey, whose population growth depends on the local basal resource state of the patch. Some fraction of predators disperse to nearby patches based on the local density of both predators and prey. A fixed proportion of prey disperse to nearby patches.

We explored two versions of the model, one with a single habitat generalist prey species, *G*, and a second with two habitat specialist prey species (*S1* and *S2*). Habitat generalists and specialists can have different responses to changes in landscape structure, potentially because of their response to particular habitats that may provide basal resources at different times of the growing season. The habitat generalist prey has high population growth in habitats A and B (gray and black in Fig. 1), thus can take advantage of different basal resources in both halves of the growing season. On the other hand, habitat specialist *S1* can have high population growth only in habitat A, and habitat specialist *S2* can have high population growth only in habitat B. Therefore, habitat specialists can take advantage of basal resources that are only abundant for a portion of the growing season.

Subpopulations of generalist prey *G* in each patch exhibit density-dependent reproduction, following a Ricker model:

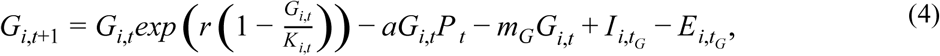

where *G*_*i,t*_ is the density of prey in patch *i* at time *t, r* is the maximum per capita rate of growth, and *I*_*i,tG*_ and *E*_*i,tG*_ represent prey immigration and emigration, respectively to and from patch *i* at time *t*. Parameter definitions and values are summarized in Table A1 (Appendix A). Prey carrying capacity *K* depends on the resource state of patch *i* at time *t*. We assumed *K* = 100 in high-resource patch states (*x*_*i*_ = 1) and *K*= 10 in low-resource patch states, (*x*_*i*_ = 0). Prey have both a constant background mortality at rate *m*_*G*_, and mortality from predation in each patch based on a per capita attack rate *a*. For the set of analyses involving two specialist prey (*S1* and *S2*), we also used equation 4 to model their population dynamics, substituting *S1* or *S2* for *G*. For simplicity we assumed a type I functional response for both generalist and specialist prey, but also explored cases where predation followed a type II functional response (Holling 1959) and found qualitatively similar results (not shown). The initial density of *G, S1*, and *S2* in each patch was set at 1. Predator and prey subpopulations that fell below a density of 1 × 10^−6^ are assumed to be locally extinct but can be recolonized by immigration from another patch.

Predator reproduction depends on the number of prey captured and the per capita conversion rate *c* of prey to predators:

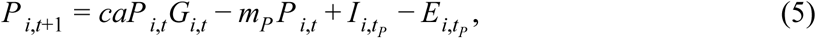

where *P*_*i,t*_ is the density of predators in patch *i* at time *t*, and *I* and *E* represent predator immigration and emigration, respectively to and from patch *i* at time *t*. Predators suffer constant mortality at rate *m*_*P*_. We used a similar equation to describe predator subpopulation dynamics in models involving two specialist prey:

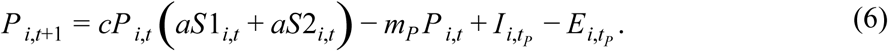

Predators and prey each have two generations in each half of the growing season (corresponding to landscape states A and B), resulting in a total of four generations per growing season. The initial density of *P* in each patch was set at 1. Predator and prey subpopulations that fell below a density of 1 × 10^−6^ are assumed to be locally extinct but can be recolonized by immigration from another patch.

Prey dispersal occurs once per generation in fixed proportion (*d*_*G*_ = 0.2) to their subpopulation size. For each patch and time step, *d*_*G*_*G*_*i,t*_ (or *d*_*S1*_*S1*_*i,t*_ or *d*_*S2*_*S2*_*i,t*_) prey move in a randomly selected direction to a new patch. Dispersal distance is randomly drawn from a Poisson distribution (*λ* = 3). Thus, prey dispersal distance is usually within one to three patches, with larger distances increasingly less likely. To avoid edge effects, the model assumes a periodic boundary condition so that dispersers that move off the landscape in one direction emerge on the opposite side. Predator dispersal follows similar rules but the fraction of each predator subpopulation dispersing *d*_*P*_ is dependent on both predator and prey density. That is, *d*_*P*_ = *µ*_*G*_ + *µ*_*P*_, where

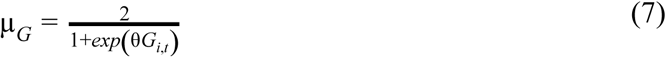

and

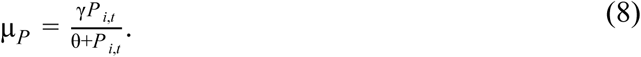

θ determines how quickly *µ*_*G*_ and *µ*_*P*_ approach their maximum and *γ* is the maximum rate of predator dispersal in response to predator density (i.e., maximum µ_2_). Therefore, *d*_*P*_ approaches 1.0 as *G*_*i,t*_ approaches 0 and *d*_*P*_ approaches *γ* as *P*_*t*_ increases. For models involving two specialist prey, *d*_*P*_ = *µ*_*S1S2*_ + *µ*_*P*_, where

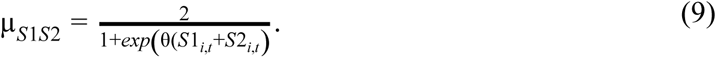

### Model analysis

Each timestep in the simulations represents a generation, thus each simulation represents 125 growing seasons (500 time steps / 4 generations per growing season). Tests showed that metapopulations always reached a dynamic equilibrium before 300 generations. Predator and prey metapopulation sizes in each timestep were quantified by summing densities across all subpopulations in a given landscape. We estimated total landscape-scale abundances of predators and prey (i.e., metapopulation sizes) as the sum of their respective abundances in each patch averaged over the final 100 timesteps of each simulation. We analyzed the combined abundance of the two specialist prey species, *S*, by summing their equilibrium metapopulation sizes (*S1* + *S2*).

We examined the dynamics of predator (*P*) and prey (*G* or *S*) metapopulations in replicate model landscapes that spanned simultaneous gradients in the season-long amount of resources (*R*) and the temporal variance in resource amount (*CV*_*R*_). Because season-long resource amount had a strong effect on metapopulation densities (described below), we examined the effects of temporal variance in each of 100 landscapes with *R* fixed at 51, 102, 154, 205, and 256 (i.e., high-resource habitat in 20, 40, 60, 80, and 100% of patches). We evaluated *R* effects on metapopulations using the proportion of maximum *R* (p*R*_max_), which for the 256-patch landscapes is *R*/256. Because predators respond directly to prey dynamics, rather than variability in *R*, we also calculated the temporal variance in prey metapopulation size (*CV*_*prey*_) over the final 100 time steps:

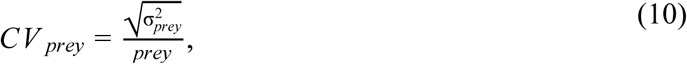

where *prey* is the equilibrium density of either *G* or *S*. We explored a range of model parameter values +/- 20% of those given in table A1, which yielded qualitatively similar results (not shown).

## Results

### Effects of resource amount and continuity on prey

Greater season-long amount of basal resources in a landscape (*R*) generally benefited prey populations. For a given level of basal resource temporal variance (*CV*_*R*_), greater p*R*_max_ increased the equilibrium abundance of generalist (*G*) and specialist prey (*S*; Fig. 2A and B). However, the effect of increased *CV*_*R*_ on prey abundance depended on prey specialization. The effect of *CV*_*R*_ on prey abundance increased with greater p*R*_max_. At higher levels of p*R*_max_, generalist prey abundance increased as *CV*_*R*_ decreased (i.e., with greater resource continuity; 2C). The opposite relationship was true for specialist prey, whose abundance increased with greater *CV*_*R*_ at higher levels of p*R*_max_ (Fig. 2D).

**Figure 2.**
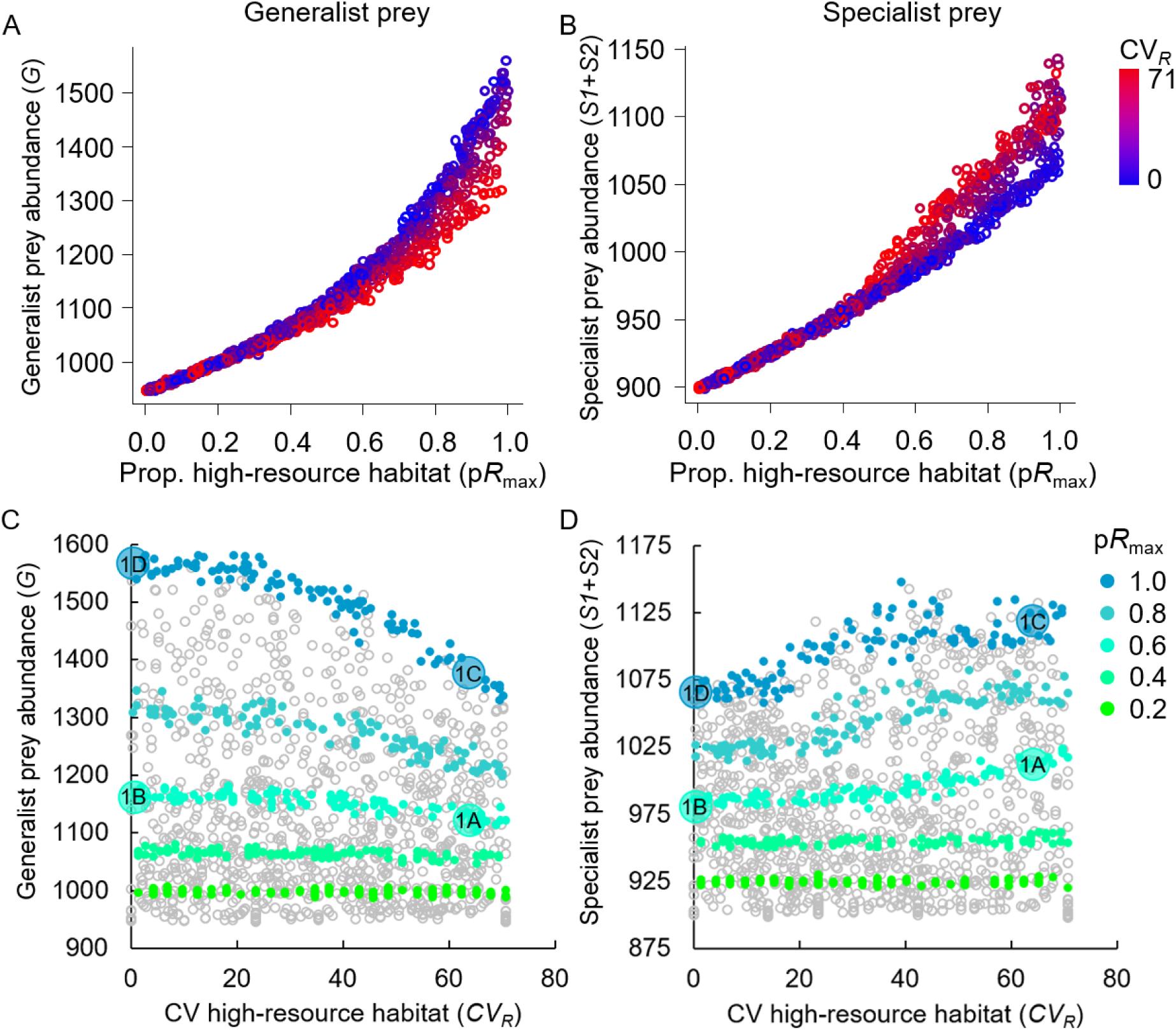
Predator and prey response to the proportion of maximum high-resource habitat in the landscape (p*R*_max_). Top panels show the density response of a generalist prey, *G* (A) and its predator, *P* (B). Bottom panels show the sum of the density response of two specialist prey, *S1+S2* (C) and their predator (D). Points represent individual model runs with landscapes varying from low (blue) to high (red) temporal variance (*CV*_*R*_).

The temporal variability of both generalist and specialist prey population (*CV*_*prey*_) increased with p*R*_max_, even when there was no temporal variability in basal resources (i.e. *CV*_*R*_ = 0, Fig. 3, blue points). This inherent effect of basal resource amount on variability in prey abundance over time was exacerbated with an increase in *CV*_*R*_ (Fig. 3, red points).

**Figure 3.**
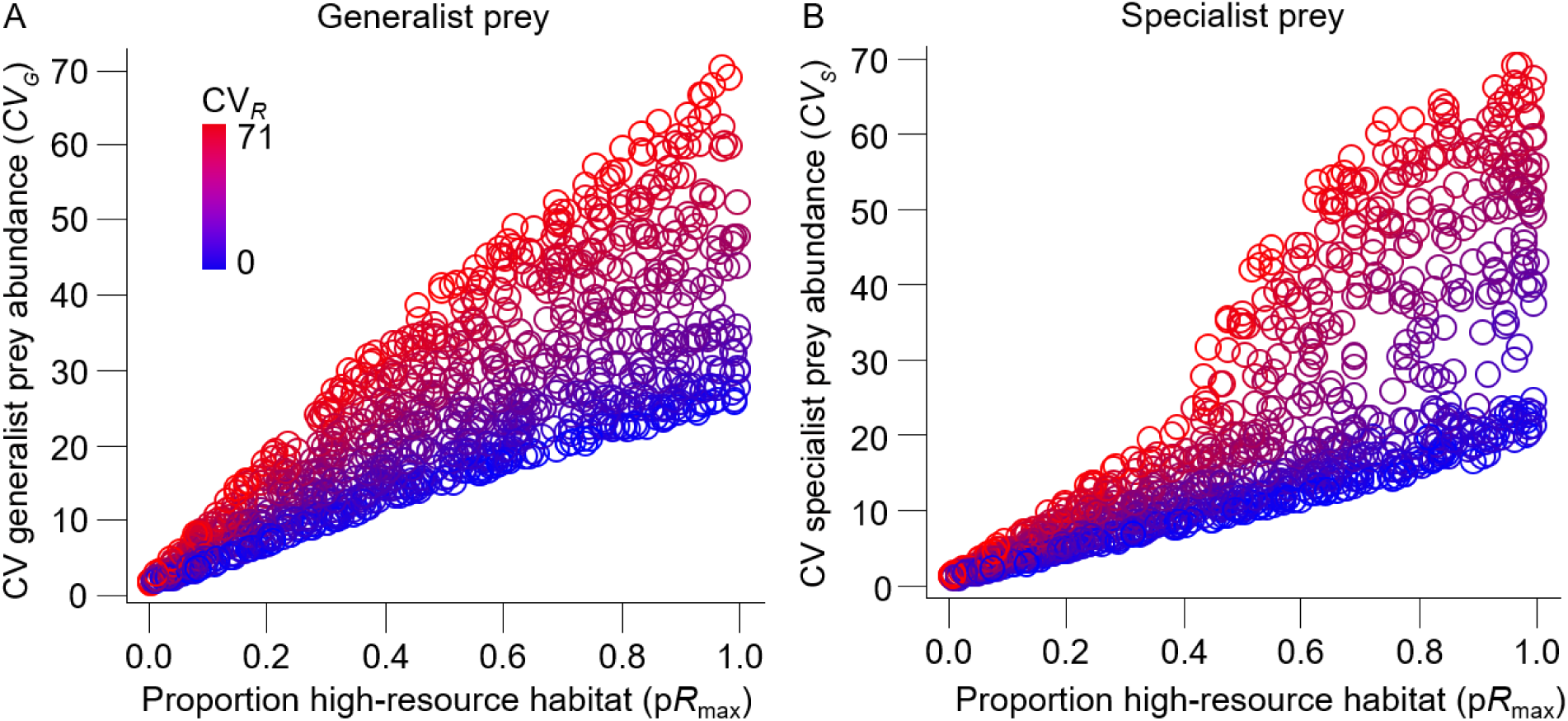
Effect of temporal variance on predator and prey density. Top panels (A, B) are model runs with one generalist prey and bottom panels (C, D) are model runs with two specialist prey. Labels within each panel (i.e., circles “1A” – “1D”) indicate model runs corresponding to the landscapes shown in Fig. 1. Colors indicate the proportion of maximum *R* (p*R*_max_), or the amount of the landscape that can provide high-resource habitat, for each of 100 separate model runs. Gray points represent a set of 1,000 model runs where p*R*_max_ was randomly selected between 0 and 1.

### Effects of resource amount and continuity on predators

The response of predators to p*R*_max_ and *CV*_*R*_ was dependent on whether prey were generalists or specialists. When feeding on generalist prey, equilibrium predator abundance counterintuitively decreased with greater p*R*_max_ across the entire range of *CV*_*R*_ (Fig. 4A). On the other hand, when feeding on specialist prey, predator abundance increased with greater p*R*_max_, but only when *CV*_*R*_ was low (i.e., in landscapes with high resource continuity; Fig. 4B). At higher levels of *CV*_*R*_, there was a non-linear relationship between predator abundance and p*R*_max_ where predator abundance initially increased with p*R*_max_ (i.e., up to p*R*_max_ ≈ 0.4) but then decreased.

**Figure 4.**
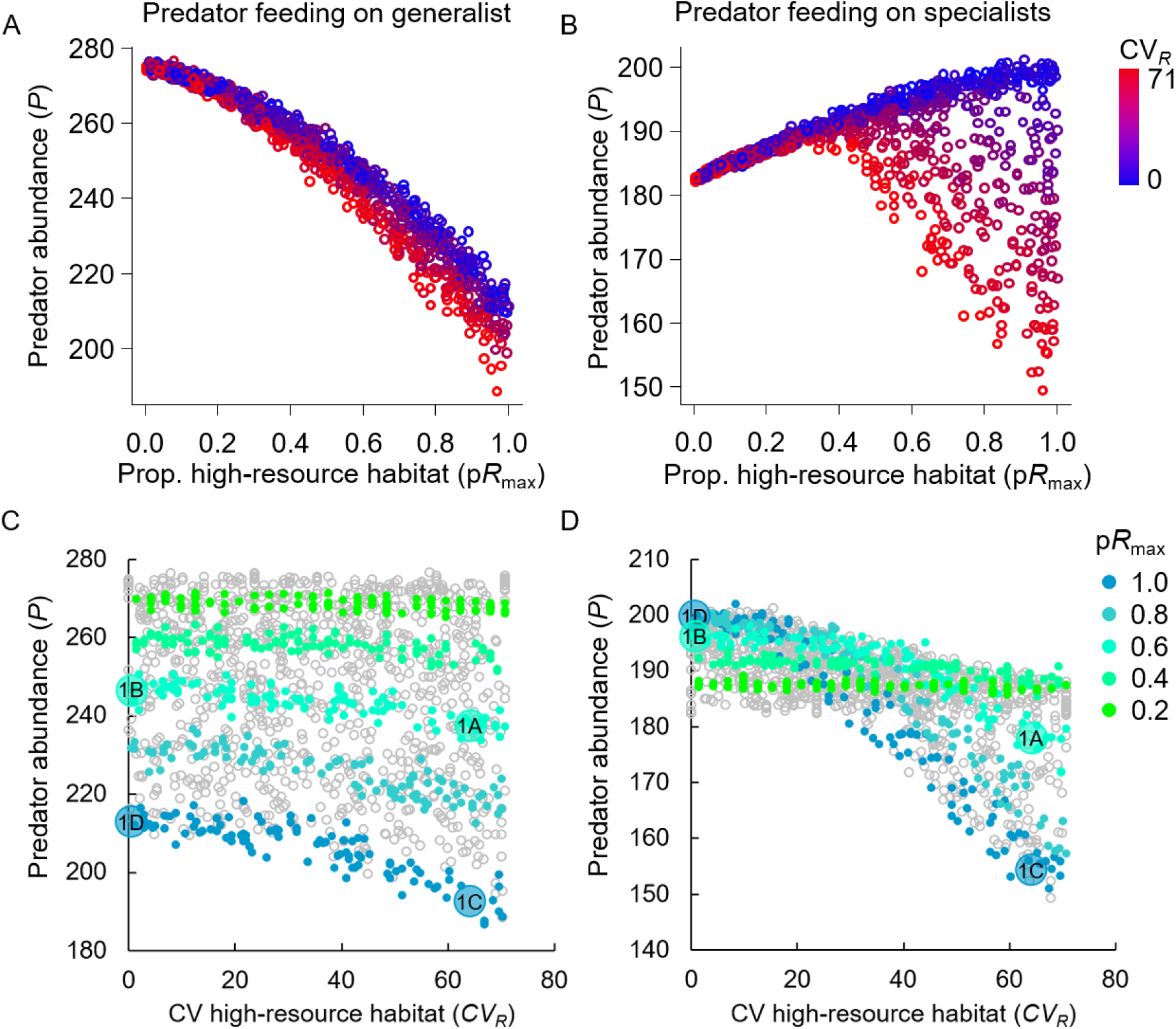
Variability in generalist (A) and total specialist (C) prey density increases with increasing proportion of maximum *R* (p*R*_max_) and with greater temporal variance in *R* (CV_*R*_, color-coded blue to red). Predator response to within-season variability in the density of generalists (B) and specialists (D) over the final 100 timesteps of each model run. Labels (1A-D) within panels B and D indicate model runs corresponding to the landscapes shown in Fig. 1. Colors in panels B and D correspond to p*R*_max_ for each of 100 separate model runs. Gray points represent a set of 1000 model runs where p*R*_max_ was randomly selected between 0 and 1.

Predator populations generally decreased with greater basal resource discontinuity (i.e., greater *CV*_*R*_), however the effect was much stronger in high-resource landscapes, especially when feeding on specialist prey (Fig. 4C and D). Predators were more sensitive to prey discontinuity, decreasing with higher *CV*_*prey*_ across all levels of p*R*_max_ (Fig. 5). As the landscape filled with more available high-resource patches (greater p*R*_max_), and/or when basal resources were available less continuously (greater *CV*_*R*_), prey populations cycled with greater amplitude, creating conditions that reduced predator populations. This is evident in example metapopulation dynamics illustrating how both p*R*_max_ and *CV*_*R*_ increase *CV*_*prey*_, and thus decrease predator abundance (Figs. B2 and B3).

**Figure 5.**
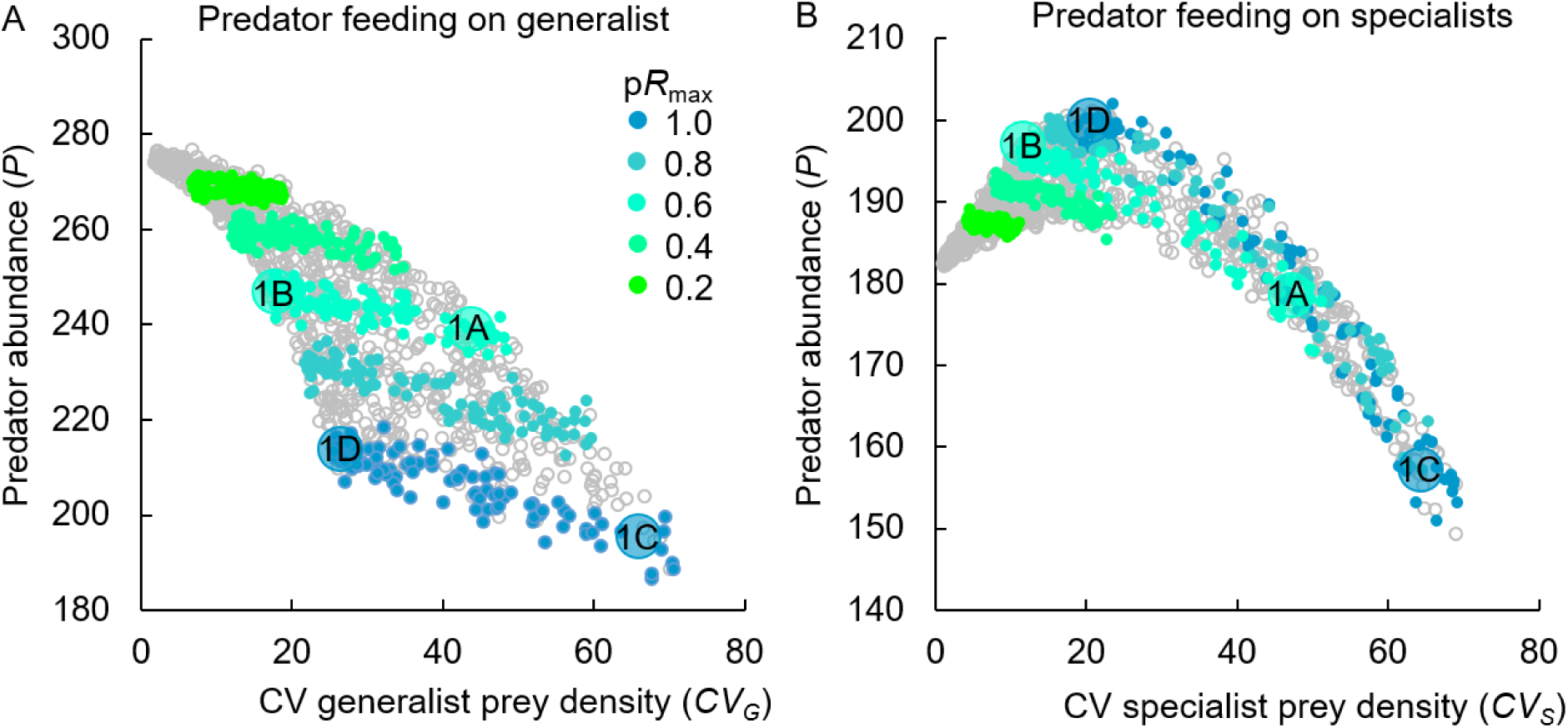
Relationship between predator and prey density for (A) one generalist prey and (B) two specialist prey. Labels (circles “1A”-“1D”) within each panel indicates model runs corresponding to the landscapes shown in Fig. 1. Colors indicate the proportion of maximum *R* (p*R*_max_), or the amount of the landscape that can provide high-resource habitat, for each of 100 separate model runs. Gray points represent a set of 1000 model runs where p*R*_max_ was randomly selected between 0 and 1.

### Prey suppression

The ability of predators to suppress prey varied with prey type, *CV*_*R*_, and p*R*_max_. In landscapes with a mid-to high-level of basal resources, lower *CV*_*R*_ (greater resource continuity) had a positive effect on both generalist prey (Fig. 2C) and their predators (Fig. 4C). Thus, for a given level of p*R*_max_, generalist prey abundance consistently increased with greater predator abundance (Fig. 6A). Furthermore, greater p*R*_max_ increased prey abundance (Fig. 3) but generally lowered predator abundance (Fig. 4A and B). Together, this suggests relatively low prey suppression by predators. That is, generalist prey populations are enhanced in low *CV*_*R*_ landscapes even though low *CV*_*R*_ also increases predator abundance. On the other hand, lower *CV*_*R*_ had a negative effect on specialist prey abundance (Fig. 2D) but a positive effect on their predators (Fig. 4D).

**Figure 6.**
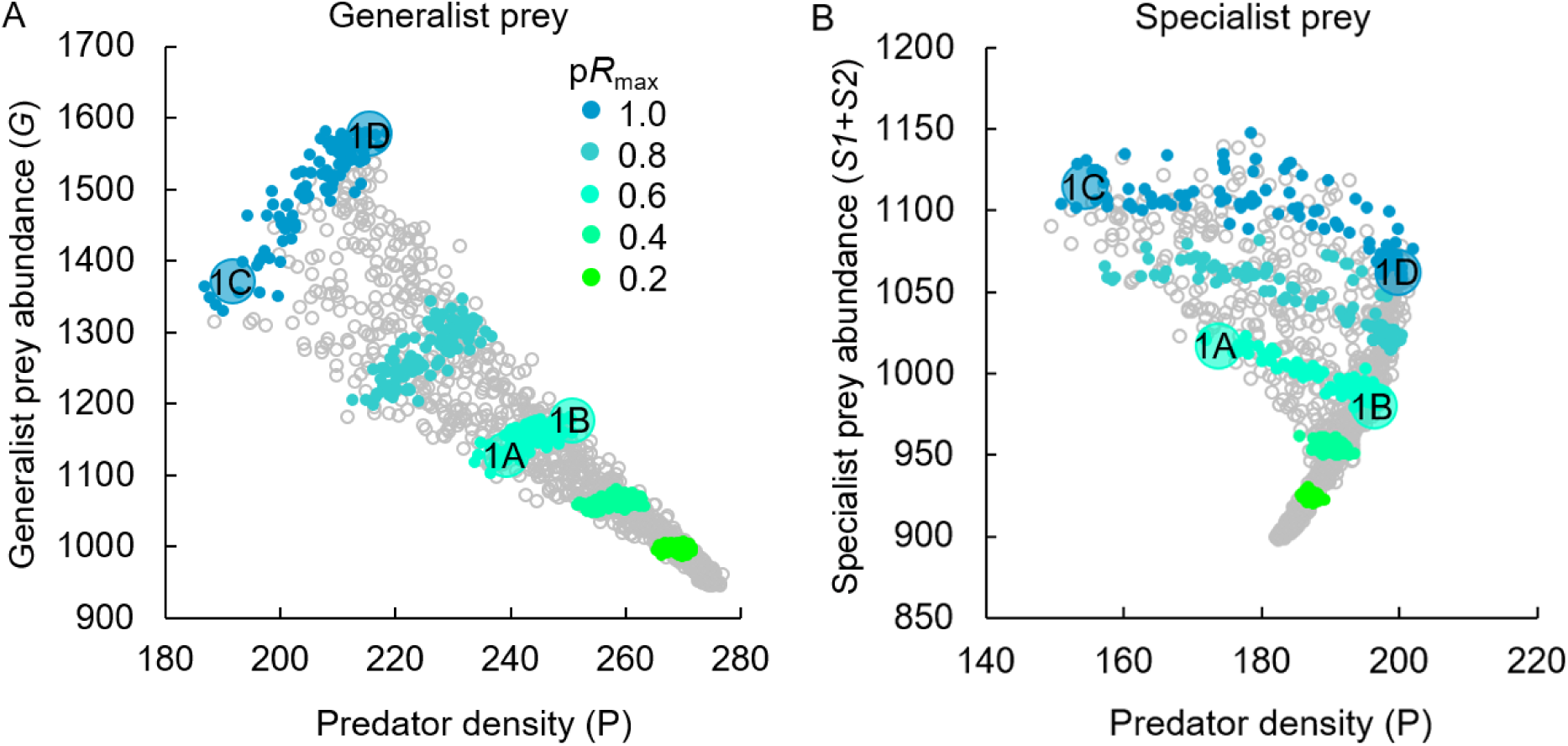
Relationship between predator and prey density for (A) one generalist prey and (B) two specialist prey. Labels (circles “1A” – “1D”) within each panel indicate model runs corresponding to the landscapes shown in Fig. 1. Colors correspond to the proportion of maximum R (pRmax), or the amount of the landscape that can provide high-resource habitat, for each of 100 separate model runs. Gray points represent a set of 1000 model runs where p*R*_max_ was randomly selected between 0 and 1.

Accordingly, specialist prey generally decreased with increasing predator abundance abundance across all levels of p*R*_max_, and especially for values of p*R*_max_ > 0.4 (Fig. 6B). This suggests relatively high prey suppression by predators. That is, when low *CV*_*R*_ increases predator populations, specialist prey declines.

## Discussion

Recent conceptual models have highlighted the importance of incorporating an understanding of the temporal dynamics of resources in a landscape, in addition to total resource amounts, in order to better understand the abundance of organisms (Schellhorn et al. 2015). Our metapopulation model showed that both the total amount of basal resources in a landscape and the temporal variance of those resources throughout a season can have large and interacting effects on population dynamics of predators and their prey. Altogether results suggest that predator abundance, and thus the potential for conservation biological control, is indirectly dependent on the temporal continuity of basal resources in the landscape and not merely the total amount. Furthermore, predator metapopulation dynamics depended on whether the prey were generalists or specialists.

### Generalist prey

A reduction in the temporal variance of resources at the landscape scale increased resource continuity for both generalist prey and their predators, and thus had a positive effect on abundance. Because generalist prey were able to use both early and late season habitats, dispersers had the opportunity to minimize the negative effects of local transition to a low-resource state and disperse to a patch potentially transitioning into a high-resource state. This temporal complementation allowed generalist prey to maintain higher metapopulation sizes even though there were also more predators in the landscape. However, the benefit of low temporal variance was most pronounced in high-resource landscapes. Low patch connectivity in low-resource landscapes may contribute to this interacting effect: when there were few high-resource patches, dispersal to another one was unlikely even if high-resource patches were present in both halves of the season.

Predators were similarly able to take advantage of a more consistent resource base by dispersing midseason to new and potentially high-resource patches. However, predator abundance declined with increasing basal resource amount and prey abundance). This type of paradox of enrichment (Rosenzweig 1971) occurs because prey populations become more temporally variable with increasing basal resource amount. The resulting fluctuations in prey abundance negatively affected predator abundance, outweighing any positive effect of more prey. Although often found in models, the paradox of enrichment is infrequently observed in natural predator-prey systems (Jensen & Ginzburg 2005). This suggests that more empirical research is necessary to explicitly assess the interacting effects of temporal variability and resource amount for predator-prey interactions in agricultural landscapes.

### Specialist prey

On the other hand, specialist prey did not benefit from lower temporal variance because they were, by design, unable to effectively use alternative resource patches when their preferred resources declined midseason. This meant that specialists could not begin a new generation in a high-resource patch at high abundance through colonization from a patch that was previously in a high-resource state. Thus, greater temporal variance actually *increased* the combined equilibrium abundance of two specialist prey species, especially in landscapes high in basal resources (Figure 2C); this is a result of increased predation in low-variance landscapes. Ives and Settle (1997) point out that synchronous crop plantings have been promoted to reduce pest outbreaks in tropical rice systems by creating temporal resource gaps for pests, but that this strategy may work only in the absence of predators. Their model agrees with ours in that in the presence of predators, asynchronous planting (i.e., low-variance landscapes) result in the lowest prey abundance.

When feeding on specialist prey, predators in our model increased in abundance with a greater proportion of high-resource patches in the landscape until about 40%, after which point predator abundance continued to increase only in landscapes with low temporal variance. In these high-resource, low-variance landscapes, the likelihood of resource gaps was sufficiently low such that predators could take advantage of greater abundance of food (prey) without populations becoming destabilized. The heterogeneity of resource availability in low-variance landscapes allows predators to maintain relatively high abundances and move into a new patch at densities capable of suppressing prey and preventing rapid prey population growth (Settle et al. 1996). However, in high-resource, high-variance landscapes, prey temporal variance was too great to maintain large predator populations.

### Applications to conservation and biological control

Increased temporal resource continuity benefits predators but also generalist prey, a potentially undesirable outcome if prey are pest species in an agricultural setting. However, this situation could be advantageous if a generalist prey species is instead the target of conservation and a desired outcome is to enhance the populations of both prey and predators (e.g., Bercovitch 2018, Roemer & Wayne 2003), if prey are not considered pests (e.g., Harwood & Obrycki 2005, Settle et al. 1996), or if the goal is biodiversity conservation in general (e.g., Harvey et al. 2020).

On the other hand, when prey are habitat specialists, greater basal resource continuity could enhance generalist predator populations at the expense of prey. Less temporally variable basal resources resulted in higher predator and lower specialist prey abundances across all levels of total habitat in the landscape. This suggests increased prey suppression (e.g., biological control of pests) in such environments. Nevertheless, the strength of this effect was diminished in landscapes with scarce basal resources, suggesting that enough high-resource patches must be present (p*R*_max_ > 0.4) for temporal variance to be relevant to biological control. A potential reason for this is that as high-resource patches become increasingly isolated in sparse landscapes, predator dispersal to a high-resource patch becomes increasingly less likely. The key feature identified by our model is that lower temporal variance benefitted predators, but only in high-resource landscapes, indicating an interaction between the temporal and resource amount aspects of a landscape.

### Limitations and future directions

This study streamlines real-world situations by making multiple simplifying assumptions. For example, the assemblage of plants, herbivores, and natural enemies in real agricultural landscapes is large, overlapping, and complex (Vandermeer et al. 2019, Snyder 2019). Future iterations of the model could vary the permutations of generalist and specialist prey and predators in the landscape. Including a mixture of generalists and specialist prey, for example, may allow for higher predator density and enhance predators’ ability to consume specialists. This could be especially true when resource variability is relatively high. Similarly, vegetation phenologies do not always separate as neatly into early- and late-season resource states as we modelhere. Thus, additional insights could be gained from new versions of this model that incorporate more complex and diverse patterns of resource phenology in the landscape.

Additionally, our model landscapes did not include habitats that provide locally continuous resources throughout the entire growing season. That is, all high-quality patch states either occurred early, or late, but not both, as may be the case in many agricultural regions. An alternative way of creating landscape-scale resource continuity for mobile consumers would be to retain habitats that provide resources throughout the entire season. These may include perennial crops (e.g., agroforestry systems, perennial grasslands, or pastures), or natural habitats (e.g. forests, prairies, or savannahs). The presence of these habitats would lower temporal variance, which, based on our model results, should increase predator abundance and in some cases prey suppression. The greater consistency of resources in such landscapes may contribute to the positive relationships often documented among non-crop area around farm fields, natural enemy populations, and biological control outcomes (Dainese et al. 2019, Chaplin-Kramer et al. 2011).

## Conclusions

The results of this modeling study support the general hypothesis that temporal discontinuity of resources in a landscape in which mobile consumers forage has a negative effect on their populations (Schellhorn et al. 2015). Landscapes with a diverse array of crop and non-crop habitats may provide resources at complementary times of the growing season, which can bolster predator populations and, in some cases, natural pest control. Indeed, recent empirical work on landscape-scale crop heterogeneity has found correlations with biological control (Redlich et al. 2018, Bosem-Baillod et al., 2017) as well as biodiversity generally across multiple trophic levels (Sirami et al. 2019). Our model provides evidence for one possible mechanism driving these relationships, namely temporal resource complementation. These results suggest that through judicious management of habitats in a landscape, either by selecting temporally complementary crops or by managing them for greater asynchrony (e.g., through planned harvest regimes), it is theoretically possible to increase predator population sizes. Furthermore, our model suggests that in addition to focusing on “habitat amount,” explicitly considering temporal dynamics will be key to understanding how landscape composition influences biodiversity and the ecosystem services.

## Acknowledgements

Support for this research came in part from USDA Grant 2018-67013-28060, the DOE Great Lakes Bioenergy Research Center (DOE BER Office of Science DE-FC02-07ER64494) and DOE OBP Office of Energy Efficiency and Renewable Energy (DE-AC05-76RL01830). We are grateful to Tania Kim, David Hoekman, and Tony Ives for feedback on earlier manuscript drafts.

## Appendices

### Appendix A: Model details

**Table A1.**
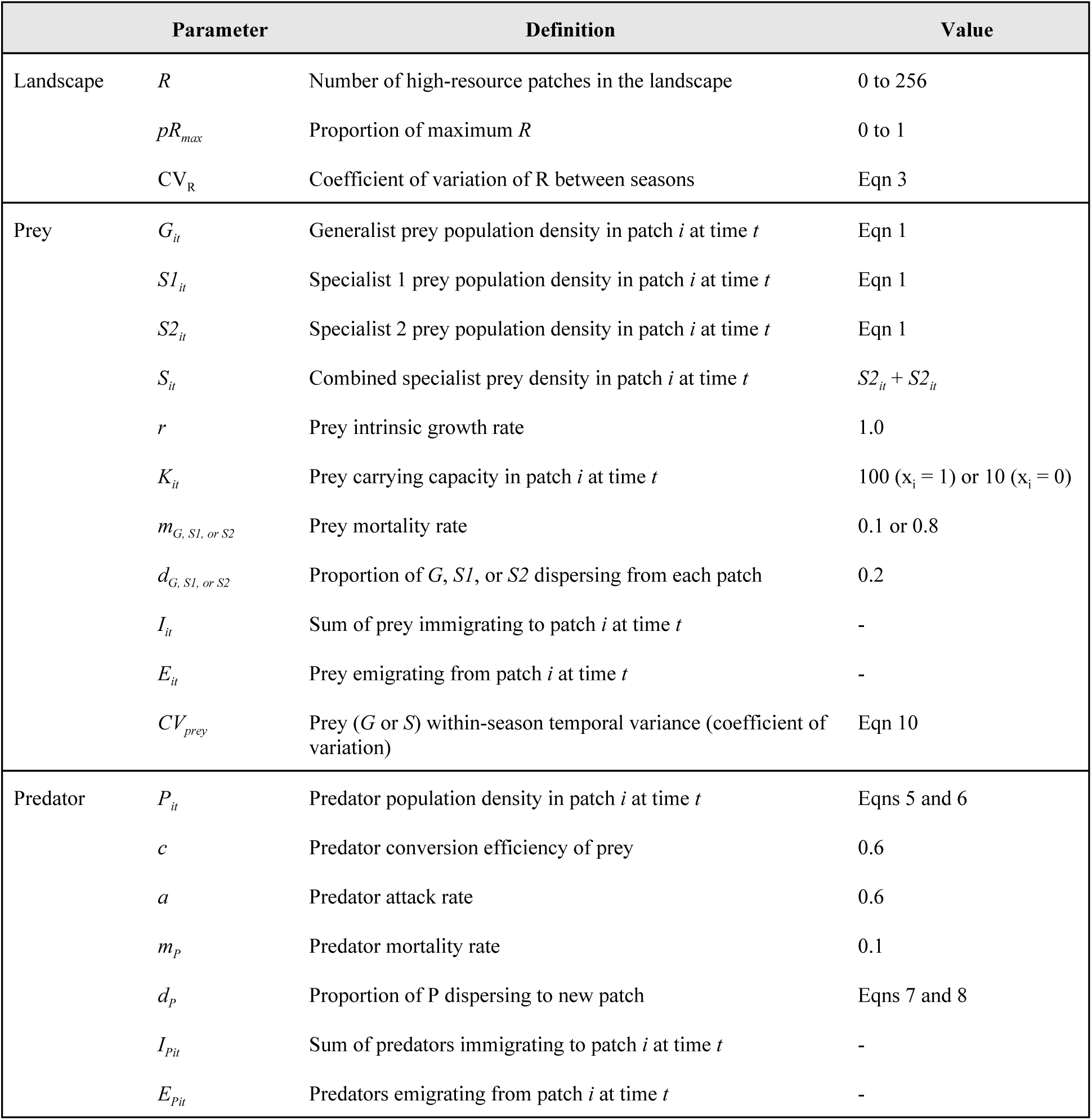
Model parameter values.

### Appendix B: Supplementary figures

**Figure B1.**
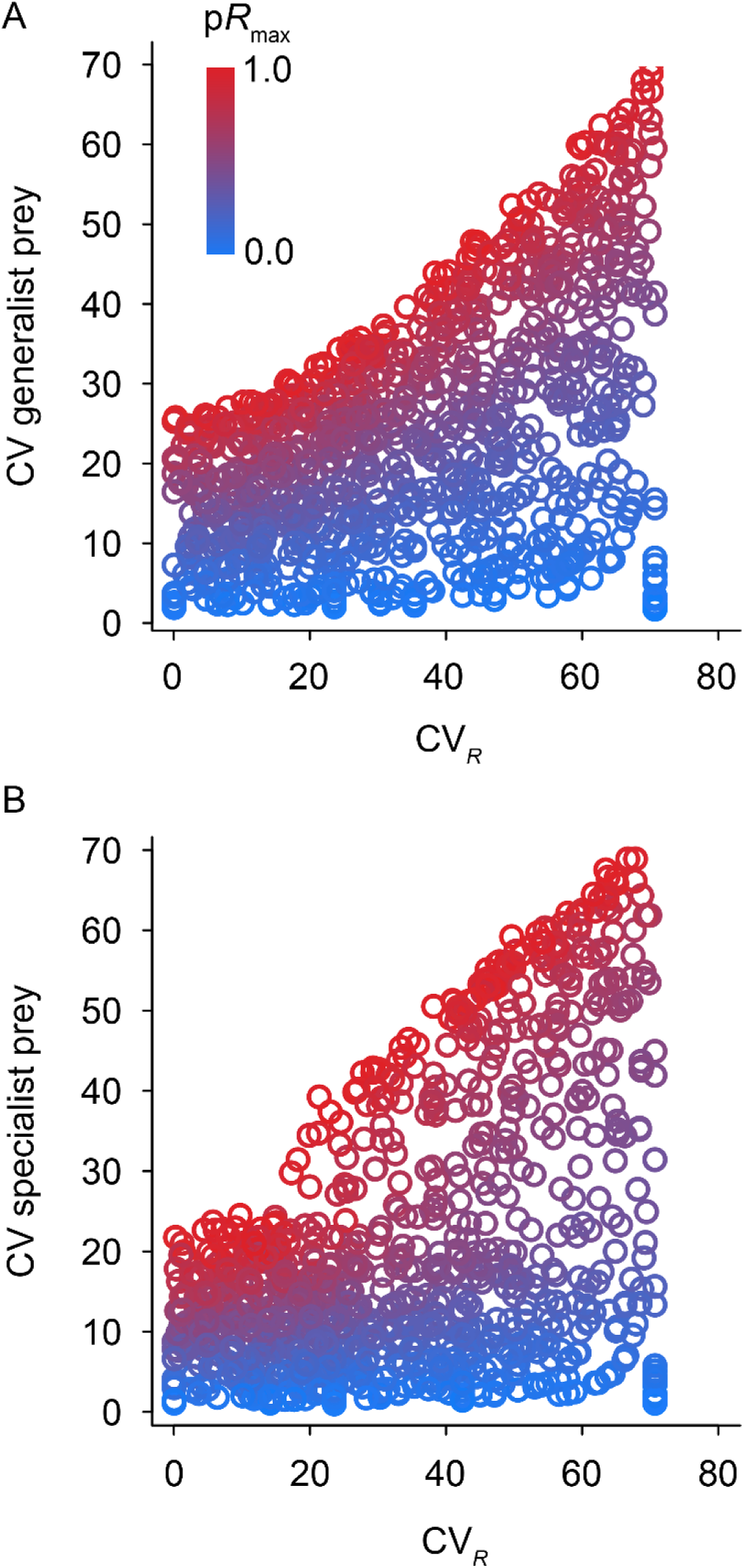
Relationship between the temporal variance of resources in landscapes and temporal variance in prey density for (A) generalists and (B) specialists. Points represent individual model runs with landscapes varying from low (blue) to high (red) p*R*_max_.

**Figure B2.**
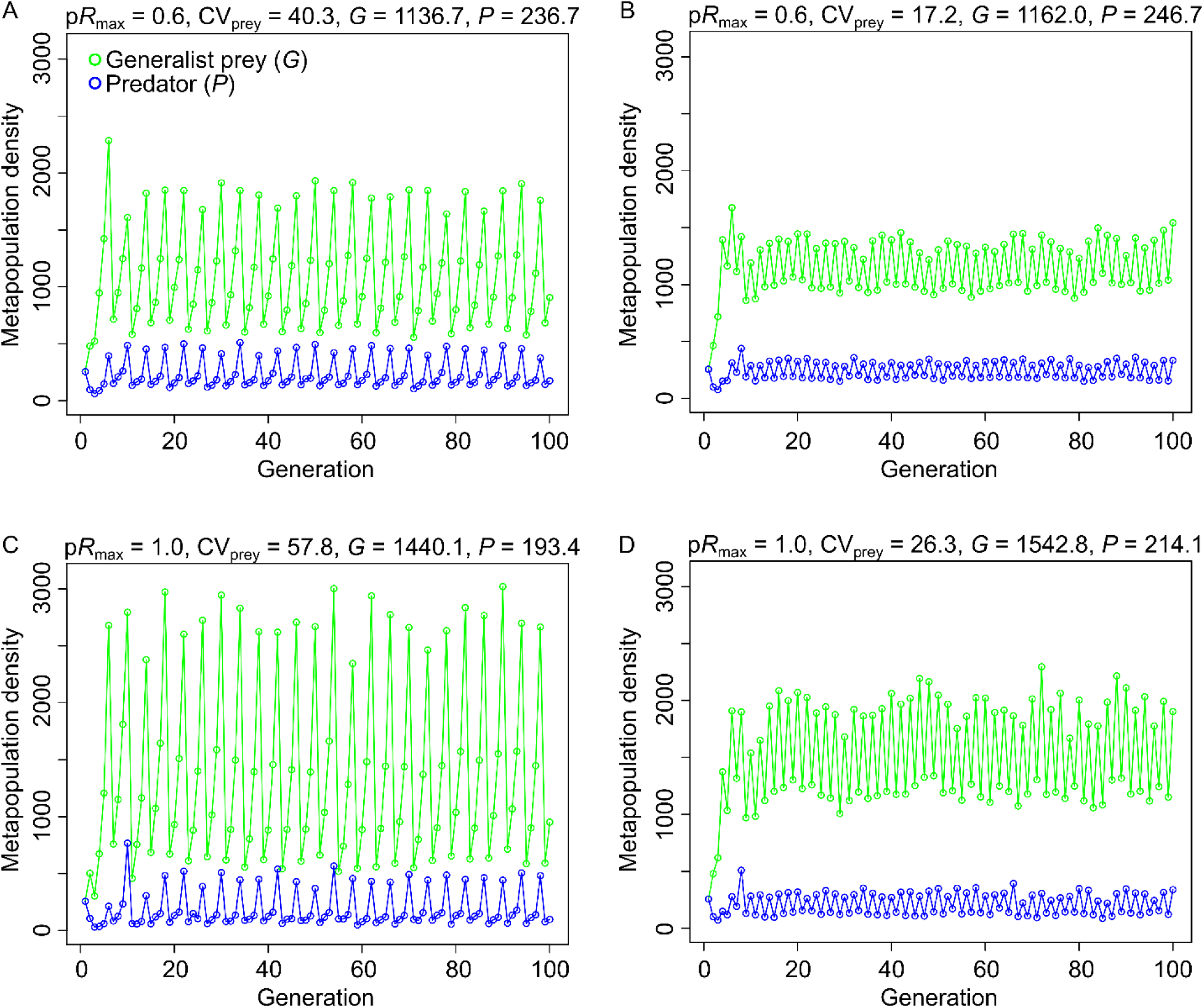
Example metapopulation dynamics of the generalist prey (*G*; green) and predator (*P*; blue). Landscapes modeled in panels A-D and associated values of *R* and *CV*_*R*_ correspond to those in Fig. 1A-D. For clarity, only the first 100 of 500 generations are shown.

**Figure B3.**
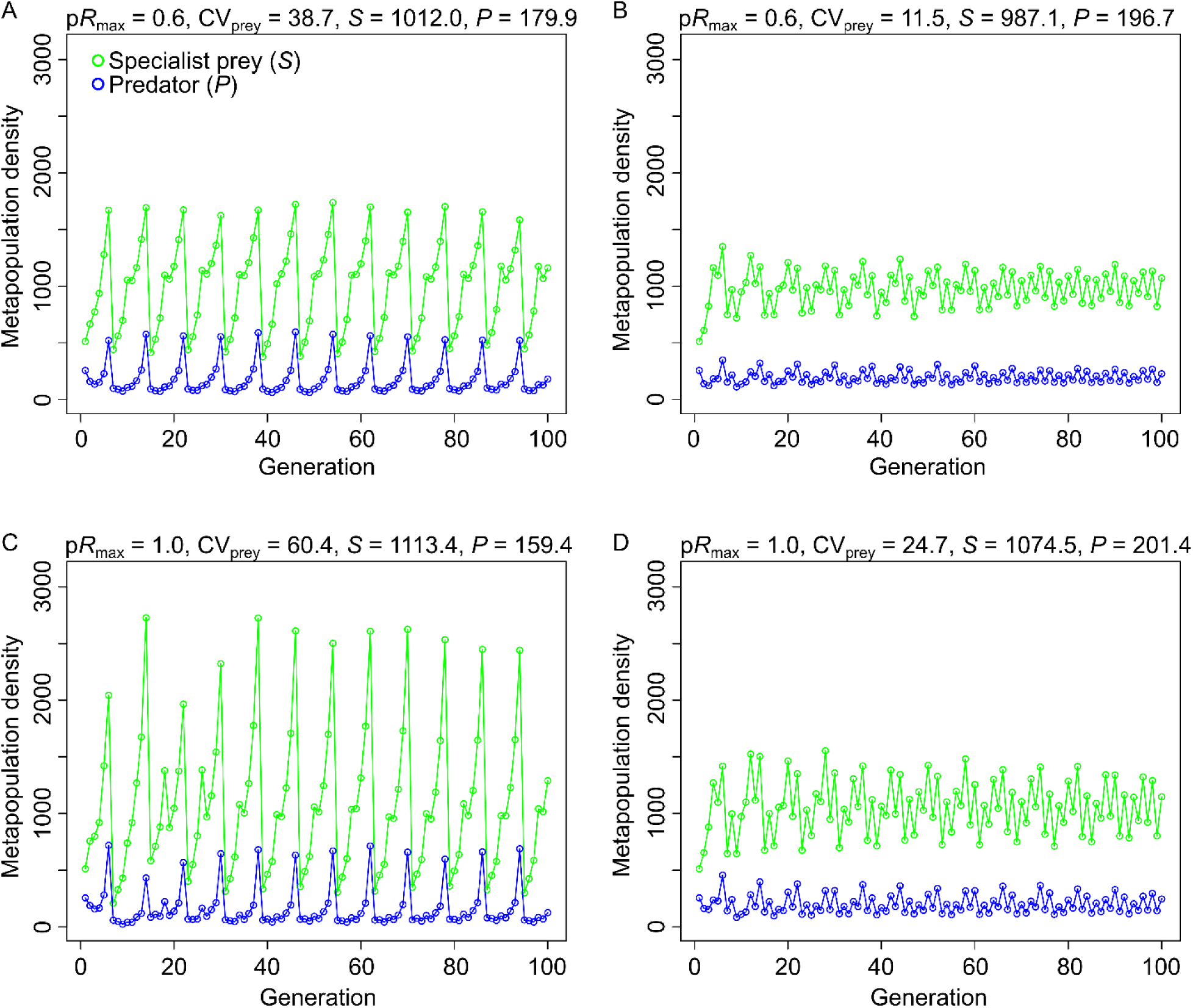
Example metapopulation dynamics of the specialist prey (*G*; green) and predator (*P*; blue). Landscapes modeled in panels A-D and associated values of *R* and *CV*_*R*_ correspond to those in Fig. 1A-D. For clarity, only the first 100 of 500 generations are shown.

