## Supplementary material for "Temporal resource continuity increases predator abundance in a metapopulation model: insights for conservation and biocontrol": R code: landscape pred prey - generalist 6.28.2019.R

```
#Brian J. Spiesman
#Ben Iuliano
#Claudio Gratton

#28 June 2019
#Metapopulation predator-prey model with a single generalist prey


rep = 100           #number of replicatate landscapes
T = 500							#number of iterations (generations) within each rep
E = 100							#number of final timesteps to calculate equilibrium abundances
RC = 16							#Number rows and columns in square matrix

disp = .2           #proportion of prey population dispersing from each cell in each timestep
dispPdmax = 0.2			#maximum predator dispersal rate based on predator density
d_p = .05						#predator dispersal rate based on predator density
d_n = .5						#predator dispersal rate based on prey density

Kgood = 100					#prey K in "good" habitat
Kbad = 10						#prey K in "bad" habitat
m = 0.1							#prey mortality rate
Mbad = .8						#prey mortality rate after change to "bad" habitat

g = 1.0             #prey intrinsic growth rate
b = 0.6							#predator conversion efficiency
a = 0.6							#predator attack rate
h = 0#0.05					#predator handling time
mP = .1							#predator mortality

disperse_i <-function(move_i, i){        #Dispersal function in i direction
	move <- i + move_i
		if(move > RC) {move = move - RC}
		if(move < 1)  {move = move + RC}
	move
}

disperse_j <-function(move_j, j){       #Dispersal function in j direction
	move <- j + move_j
		if(move > RC) {move = move - RC}
		if(move < 1)  {move = move + RC}
	move
}

#Placeholders for summary results
Nabund = matrix(0,rep,1)
Pabund = matrix(0,rep,1)
HabA = matrix(0,rep,1)
HabB = matrix(0,rep,1)
VarHab = matrix(0,rep,1)
TotHab = matrix(0,rep,1)
VarLandN = matrix(0,rep,1)

	pb = winProgressBar(title = "progress bar", min = 0, max = rep, width = 300)				        #Setup progress bar
for (r in 1:rep){ #Begin looping through landscape replicates
	Sys.sleep(0.0001);setWinProgressBar(pb, r, title=paste(round(r/rep*100, 1),"% complete"))		#Start progress bar

  #Set up landscape with total seasonl long "good" good habitat locations
	H = runif(1)                    #Proportion of "good" habitat present in the landscape
	Habitat = round((RC*RC)*H)			#Number of matrix entries that can be "good"
	CLands = matrix(0,RC,RC)				#Matrix of "bad" (0) and "good" (1) locations
	while(sum(CLands) < Habitat){
		x = round(runif(1,1,(RC*RC)))
		CLands[x] = 1
	}

	#Divide the "good" habitat between the first and second halves of the season 
	H1 = H*runif(1,0,1) 						#Proportion of "good" habitat present in the first half of season
	Habitat1 = round(RC*RC*H1)			#Number of "good" matrix entries in the first half of the season
	Lands1 = matrix(0,RC,RC)				#Landscape state with good (1) and bad (0) habitat during the first half of the season
	while(sum(Lands1) < Habitat1){
		x = round(runif(1,1,(RC*RC)))
		if(CLands[x]==1){Lands1[x]=1}
	}
	Lands2 = CLands-Lands1          #Landscape state with good (1) and bad (0) habitat during the second half of the season

	N = array(1,dim=c(RC,RC,T))			#Array of prey population sizes in each landscape cell in each timestep
	P = array(1,dim=c(RC,RC,T))			#Array of predator population sizes in each landscape cell in each timestep
	
	Lands_previous = list(Lands1)

	#Population dynamics and dispersal
	for (t in 2:T){      #Changes landscape from early structure to late structure at the start of the 3rd timestep of each season (4 timesteps per season)
		change = t%%4
		if(change == 0) {change = 4}
		if(change < 3) {Lands=Lands1}	
		else{Lands = Lands2}
    #Prey mortality associated with resource change (i.e., local habitat changes from good to bad)
		lands.prev = Lands_previous[[(t-1)]]
		for (i in 1:RC){
			for (j in 1:RC){
				if(Lands[i,j] - lands.prev[i,j]==-1){N[i,j,t] = (1-Mbad)*N[i,j,(t-1)]}
				else{N[i,j,t] = N[i,j,(t-1)]}
			}
		}
		Lands_previous[[t]] = Lands #update previous landscape structure for next timestep
    #Prey dispersal
		temp_disp = matrix(0,RC,RC) 
    for (i in 1:RC){
      for (j in 1:RC){
			  move_i = rpois(1, lambda = 3)+1                 #dispersal distance sampled from poisson distribution
        move_j = rpois(1, lambda = 3)+1       #dispersal distance sampled from poisson distribution
				if(move_i > 15){move_i=15}
				if(move_j > 15){move_j=15}
          target_i = disperse_i(move_i,i)
          target_j = disperse_j(move_j,j)
          temp_disp[target_i, target_j] = ((disp*N[i,j,(t-1)]) + temp_disp[target_i,target_j])        #add number dispersing to proper cell of temp_disp matrix
          temp_disp[i,j] = (temp_disp[i,j] - (disp*N[i,j,(t-1)]))                                     #subtract individuals from the cell in which they just moved
      }
		}
		N[,,t] = temp_disp[,] + N[,,t]

    #Predator dispersal
		temp_disp = matrix(0,RC,RC) 
    for (i in 1:RC){
      for (j in 1:RC){
			  dispNd = (2/(1+exp(d_n*N[i,j,t])))							          #portion of predator dispersal rate dependent on prey density
			  dispPd = ((dispPdmax*P[i,j,(t-1)])/(d_p+P[i,j,(t-1)]))		#portion of predator dispersal rate dependent on pred density
			  dispersal = dispNd + dispPd
				move_i = rpois(1, lambda = 3)+1                           #dispersal distance sampled from poisson distribution
        move_j = rpois(1, lambda = 3)+1               	          #dispersal distance sampled from poisson distribution
				if(move_i > 15){move_i=15}
				if(move_j > 15){move_j=15}
          target_i = disperse_i(move_i,i)
          target_j = disperse_j(move_j,j)
          temp_disp[target_i, target_j] = ((dispersal*P[i,j,(t-1)]) + temp_disp[target_i,target_j])  #add number dispersing to proper cell of temp_disp matrix
          temp_disp[i,j] = (temp_disp[i,j] - (dispersal*P[i,j,(t-1)]))                               #subtract individuals from the cell in which they just moved
      }
		}
		P[,,t] = temp_disp[,] + P[,,(t-1)]

    #Pop dynamics
		for (i in 1:RC){
      for (j in 1:RC){
        if (Lands[i,j] == 1) {K = Kgood}		#K (carrying capacity) when in landscape habitat type 1
        else{K = Kbad}								    #K (carrying capacity) when in landscape habitat type 0      		
        N[i,j,t] = (N[i,j,t]*exp(g*(1-(N[i,j,t]/K))))-(a*N[i,j,t]*P[i,j,t])/(1+(a*h*N[i,j,t]*P[i,j,t])) - (m*N[i,j,t]) #ricker model
					if (N[i,j,t]<0.0000001){N[i,j,t]=0}						#trap: makes local pop size zero if below 0.0000001
				P[i,j,t] = (b*a*N[i,j,t]*P[i,j,t])/(1 + (a*h*N[i,j,t]*P[i,j,t])) - (mP*P[i,j,t])
					if (P[i,j,t]<0.0000001){P[i,j,t]=0}						#trap: makes local pop size zero if below 0.0000001
			}
		}
	}
	Nequalibabund = array(0,dim=c(RC,RC))
	for (i in 1:RC){
    for (j in 1:RC){
      Nequalibabund[i,j] = mean(N[i,j,(T-E):T])		#mean abundance in each cell over the final E timesteps
    }
	}
	Nabundmap = Nequalibabund

	Nlandabund = seq(1:T)					#total prey abundance in the landscape at each timestep
	for (t in 1:T){
		Nlandabund[t] = sum(N[,,t])
	}

	Pequalibabund = array(0,dim=c(RC,RC))
	for (i in 1:RC){
    for (j in 1:RC){
      Pequalibabund[i,j] = mean(P[i,j,(T-E):T])		#mean abundance in each cell over the final 4 timesteps
    }
	}
	Pabundmap = Pequalibabund

	Plandabund = seq(1:T)					#total predator abundance in the landscape at each timestep
	for (t in 1:T){
		Plandabund[t] = sum(P[,,t])
	}
  #Summary values
	Nabund[r] = sum(Nequalibabund)      #Total mean prey abundance
	Pabund[r] = sum(Pequalibabund)      #Total mean pred abundance
	HabA[r] = sum(Lands1)               #Amount of good habitat in the first half of the season
	HabB[r] = sum(Lands2)               #Amount of good habitat in the second half of the season
	TotHab[r] = sum(HabA[r],HabB[r])    #Season-long amount of good habitat
	VarHab[r] = var(c(HabA[r],HabB[r])) #Landscape-scale temporal variance
	VarLandN[r] = var(Nlandabund[(T-E):T])
}

close(pb)		#Turn off progress bar

CVHab = 100*sqrt(VarHab)/TotHab      #Calculate the CV of basal resource temporal variance
CVLandN = 100*sqrt(VarLandN)/Nabund  #Calculate the CV of prey temporal variance

output = cbind(seq(1:rep),Nabund,Pabund,TotHab,VarHab,CVHab,VarLandN,CVLandN,HabA,HabB)
colnames(output) = c("Run","Nabund","Pabund","TotHab","VarHab","CVHab","VarLandN","CVLandN","HabA","HabB")
output = as.data.frame(output)
output

par(mfrow=c(3,2))
plot(Nabund ~ TotHab,col="black", ylab="Total prey abund", xlab="Habitat amount")
plot(Pabund ~ TotHab,col="black", ylab="Total predator abund", xlab="Habitat amount")
plot(Nabund ~ CVHab,col="black", ylab="Total prey abund", xlab="CV habitat")
plot(Pabund ~ CVHab,col="black", ylab="Total predator abund", xlab="CV habitat")
plot(Nabund~ Pabund,col="black", ylab="Total prey abund", xlab="Total predator abund")
plot(CVHab ~ TotHab="black", ylab="CV habitat", xlab="Habitat amount")
```
