## Supplementary material for "Temporal resource continuity increases predator abundance in a metapopulation model: insights for conservation and biocontrol": R code: landscape pred prey - two specialists 6.28.2019.R

```
#Brian J. Spiesman
#Ben Iuliano
#Claudio Gratton

#28 June 2019
#Metapopulation predator-prey model with two specialist prey

rep = 100           #number of replicatate landscapes
T = 500							#number of iterations (generations) within each rep
E = 100							#number of final timesteps to calculate equilibrium abundances
RC = 16							#Number rows and columns in square matrix

disp = .2           #proportion of prey population dispersing from each cell in each timestep
dispPdmax = 0.2			#maximum predator dispersal rate based on predator density
d_p = .05						#predator dispersal rate based on predator density
d_n = .5						#predator dispersal rate based on prey density

Kgood = 100					#prey K in "good" habitat
Kbad = 10						#prey K in "bad" habitat
m = 0.1							#prey mortality rate
Mbad = .8						#prey mortality rate after change to "bad" habitat

g = 1.0             #prey intrinsic growth rate
b = 0.6							#predator conversion efficiency
a = 0.6							#predator attack rate
h = 0#0.05					#predator handling time
mP = .1							#predator mortality

disperse_i <-function(move_i, i){        #Dispersal function in i direction
  move <- i + move_i
  if(move > RC) {move = move - RC}
  if(move < 1)  {move = move + RC}
  move
}

disperse_j <-function(move_j, j){       #Dispersal function in j direction
  move <- j + move_j
  if(move > RC) {move = move - RC}
  if(move < 1)  {move = move + RC}
  move
}

#Placeholders for summary results
N1abund = matrix(0,rep,1)
N2abund = matrix(0,rep,1)
NTabund = matrix(0,rep,1)
Pabund = matrix(0,rep,1)
HabA = matrix(0,rep,1)
HabB = matrix(0,rep,1)
VarHab = matrix(0,rep,1)
TotHab = matrix(0,rep,1)
VarLandNT = matrix(0,rep,1)

	pb = winProgressBar(title = "progress bar", min = 0, max = rep, width = 300)				#Setup progress bar
for (r in 1:rep){
	Sys.sleep(0.0001);setWinProgressBar(pb, r, title=paste(round(r/rep*100, 1),"% complete"))		#Start progress bar

  #Set up landscape with total seasonl long "good" good habitat locations
  H = runif(1)                    #Proportion of "good" habitat present in the landscape
  Habitat = round((RC*RC)*H)			#Number of matrix entries that can be "good"
  CLands = matrix(0,RC,RC)				#Matrix of "bad" (0) and "good" (1) locations
  while(sum(CLands) < Habitat){
    x = round(runif(1,1,(RC*RC)))
    CLands[x] = 1
  }
  
  #Divide the "good" habitat between the first and second halves of the season 
  H1 = H*runif(1,0,1) 						#Proportion of "good" habitat present in the first half of season
  Habitat1 = round(RC*RC*H1)			#Number of "good" matrix entries in the first half of the season
  Lands1 = matrix(0,RC,RC)				#Landscape state with good (1) and bad (0) habitat during the first half of the season
  while(sum(Lands1) < Habitat1){
    x = round(runif(1,1,(RC*RC)))
    if(CLands[x]==1){Lands1[x]=1}
  }
  Lands2 = CLands-Lands1          #Landscape state with good (1) and bad (0) habitat during the second half of the season
  
  N1 = array(1,dim=c(RC,RC,T))				#Array of prey1 population sizes in each landscape cell in each time step
	N2 = array(1,dim=c(RC,RC,T))				#Array of prey2 population sizes in each landscape cell in each time step
	NT = array(1,dim=c(RC,RC,T))				#Array of total prey population sizes in each landscape cell in each time step
	P = array(1,dim=c(RC,RC,T))				  #Array of predator population sizes in each landscape cell in each time step

	Lands_previous = array(1,dim=c(RC,RC,T))
	Lands_previous[,,1] = Lands1

	
	#Population dynamics and dispersal
	for (t in 2:T){
		change = t%%4                #Changes landscape from early structure to late structure at the start of the 3rd timestep of each season (4 timesteps per season)
		if(change == 0) {change = 4}
		if(change < 3) {Lands=Lands1}	
		else{Lands = Lands2}

    #Prey mortality associated with resource change (i.e., local habitat changes from good to bad)
	  lands.prev = Lands_previous[,,(t-1)]
		for (i in 1:RC){
			for (j in 1:RC){
				if(Lands[i,j] - lands.prev[i,j]==-1){N1[i,j,t] = (1-Mbad)*N1[i,j,(t-1)]}
				else{N1[i,j,t] = N1[i,j,(t-1)]}
			}
		}
		for (i in 1:RC){
			for (j in 1:RC){
				if(Lands[i,j] - lands.prev[i,j]==-1){N2[i,j,t] = (1-Mbad)*N2[i,j,(t-1)]}
				else{N2[i,j,t] = N2[i,j,(t-1)]}
			}
		}
		Lands_previous[,,t] = Lands

    #N1 dispersal
		temp_disp = matrix(0,RC,RC) 
      	for (i in 1:RC){
          for (j in 1:RC){
				    move_i = rpois(1, lambda = 3)+1                 #dispersal distance sampled from poisson distribution
            move_j = rpois(1, lambda = 3)+1               	#dispersal distance sampled from poisson distribution
					    if(move_i > 15){move_i=15}
					    if(move_j > 15){move_j=15}
                target_i = disperse_i(move_i,i)
                target_j = disperse_j(move_j,j)
                temp_disp[target_i, target_j] = ((disp*N1[i,j,t]) + temp_disp[target_i,target_j])        #add number dispersing to proper cell of temp_disp matrix
                temp_disp[i,j] = (temp_disp[i,j] - (disp*N1[i,j,t]))                                     #subtract individuals from the cell in which they just moved
          }
		    }
		N1[,,t] = temp_disp[,] + N1[,,t]

    #N2 dispersal
		temp_disp = matrix(0,RC,RC) 
    for (i in 1:RC){
      for (j in 1:RC){
			move_i = rpois(1, lambda = 3)+1                 #dispersal distance sampled from poisson distribution
      move_j = rpois(1, lambda = 3)+1               	#dispersal distance sampled from poisson distribution
				if(move_i > 15){move_i=15}
				if(move_j > 15){move_j=15}
          target_i = disperse_i(move_i,i)
          target_j = disperse_j(move_j,j)
          temp_disp[target_i, target_j] = ((disp*N2[i,j,t]) + temp_disp[target_i,target_j])        #add number dispersing to proper cell of temp_disp matrix
          temp_disp[i,j] = (temp_disp[i,j] - (disp*N2[i,j,t]))                                     #subtract individuals from the cell in which they just moved
      }
		}
		N2[,,t] = temp_disp[,] + N2[,,t]

		NT[,,t] = N1[,,t] + N2[,,t]

		#Predator dispersal
		temp_disp = matrix(0,RC,RC) 
      for (i in 1:RC){
        for (j in 1:RC){
					dispNd = (2/(1+exp(d_n*NT[i,j,t])))						              #portion of predator dispersal rate dependent on prey density
					dispPd = ((dispPdmax*P[i,j,(t-1)])/(d_p+P[i,j,(t-1)]))			#portion of predator dispersal rate dependent on pred density
					dispersal = dispNd + dispPd
					move_i = rpois(1, lambda = 3)+1                             #dispersal distance sampled from poisson distribution
          move_j = rpois(1, lambda = 3)+1               	            #dispersal distance sampled from poisson distribution
						if(move_i > 15){move_i=15}
						if(move_j > 15){move_j=15}
              target_i = disperse_i(move_i,i)
              target_j = disperse_j(move_j,j)
              temp_disp[target_i, target_j] = ((dispersal*P[i,j,(t-1)]) + temp_disp[target_i,target_j])   #add number dispersing to proper cell of temp_disp matrix
              temp_disp[i,j] = (temp_disp[i,j] - (dispersal*P[i,j,(t-1)]))                                #subtract individuals from the cell in which they just moved	
        }
			}
			P[,,t] = temp_disp[,] + P[,,(t-1)]

    #Pop dynamics
		for (i in 1:RC){
      for (j in 1:RC){
        #N1 population dynamics
        if (Lands[i,j] == 3) {K = Kgood; m = Mgood}				#K (carrying capacity) when in landscape habitat type 1
        else{K = Kbad; m = Mbad}								          #K (carrying capacity) when in landscape habitat type 0      		
          N1[i,j,t] = (N1[i,j,t]*exp(g*(1-(N1[i,j,t]/K))))-(a*N1[i,j,t]*P[i,j,t])/(1+(a*h*N1[i,j,t]*P[i,j,t])) - (m*N1[i,j,t])
					if (N1[i,j,t]<0.0000001){N1[i,j,t]=0}						#trap - makes local pop size zero if below 0.0000001

        #N2 population dynamics
        if (Lands[i,j] == 4) {K = Kgood; m = Mgood}				#K (carrying capacity) when in landscape habitat type 1
        else{K = Kbad; m = Mbad}
          N2[i,j,t] = (N2[i,j,t]*exp(g*(1-(N2[i,j,t]/K))))-(a*N2[i,j,t]*P[i,j,t])/(1+(a*h*N2[i,j,t]*P[i,j,t])) - (m*N2[i,j,t])
				  if (N2[i,j,t]<0.0000001){N2[i,j,t]=0}					#trap - makes local pop size zero if below 0.0000001

        #Predator population dynamics
        P[i,j,t] = b*P[i,j,t] * (((a*N1[i,j,t])+(a*N2[i,j,t]))/(1 + (((a*h*N1[i,j,t])+(a*h*N2[i,j,t]))*P[i,j,t]))) - (mP*P[i,j,t]) #predator model
				if (P[i,j,t]<0.0000001){P[i,j,t]=0}						     #trap - makes local pop size zero if below 0.0000001
			}
		}
	}
	N1equalibabund = array(0,dim=c(RC,RC))
	for (i in 1:RC){
    for (j in 1:RC){
      N1equalibabund[i,j] = mean(N1[i,j,(T-E):T])		#mean abundance in each cell over the final E timesteps
    }
	}
	N2equalibabund = array(0,dim=c(RC,RC))
	for (i in 1:RC){
    for (j in 1:RC){
      N2equalibabund[i,j] = mean(N2[i,j,(T-E):T])		#mean abundance in each cell over the final E timesteps
    }
	}
	NTequalibabund = N1equalibabund + N2equalibabund
	N1landabund = seq(1:T)					#total abundance in the landscape at each timestep
	for (t in 1:T){
		N1landabund[t] = sum(N1[,,t])
	}
	N2landabund = seq(1:T)					#total abundance in the landscape at each timestep
	for (t in 1:T){
		N2landabund[t] = sum(N2[,,t])
	}
	NTlandabund = N1landabund + N2landabund
	Pequalibabund = array(0,dim=c(RC,RC))
	for (i in 1:RC){
    for (j in 1:RC){
      Pequalibabund[i,j] = mean(P[i,j,(T-E):T])		#mean abundance in each cell over the final 4 timesteps
    }
	}
	Plandabund = seq(1:T)					#total abundance in the landscape at each timestep
	for (t in 1:T){
		Plandabund[t] = sum(P[,,t])
	}
	N1abund[r] = sum(N1equalibabund)
	N2abund[r] = sum(N2equalibabund)
	NTabund[r] = sum(NTequalibabund)
	Pabund[r] = sum(Pequalibabund)
	HabA[r] = sum(Lands1)
	HabB[r] = sum(Lands2)
	TotHab[r] = sum(HabA[r],HabB[r])
	VarHab[r] = var(c(HabA[r],HabB[r]))
	VarLandNT[r] = var(NTlandabund[(T-E):T])
}
close(pb)		#Turn off progress bar

CVHab = 100*sqrt(VarHab)/TotHab         #Calculate the CV of basal resource temporal variance
CVLandNT = 100*sqrt(VarLandNT)/NTabund  #Calculate the CV of total prey temporal variance

output = cbind(seq(1:rep),N1abund,N2abund,NTabund,Pabund,TotHab,VarHab,CVHab,VarLandNT,CVLandNT,HabA,HabB)
colnames(output) = c("Run","N1abund","N2abund","NTabund","Pabund","TotHab","VarHab","CVHab","VarLandNT","CVLandNT","HabA","HabB")
output = as.data.frame(output)
output

par(mfrow=c(3,2))
plot(NTabund ~ VarHab,col="black", ylab="Total prey abund (N1+N2)", xlab="Landscape temporal variance")
plot(Pabund ~ VarHab,col="black", ylab="Total predator abund", xlab="Landscape temporal variance")
plot(Pabund ~ CVHab,col="black", ylab="Total predator abund", xlab="CV habitat")
plot(Pabund ~ CVLandNT,col="black", ylab="Total predator abund", xlab="CV prey")
plot(NTabund~ Pabund,col="black", ylab="Total prey abund (N1+N2)", xlab="Total predator abund")
plot(NTabund ~ TotHab,col="black", ylab="Total predator abund", xlab="Habitat amount")
```
